# Antiepileptic medication strengthens excitatory neurotransmission in pyramidal neurons of the adult human neocortex

**DOI:** 10.1101/2023.09.30.560289

**Authors:** Maximilian Lenz, Pia Kruse, Amelie Eichler, Jakob Straehle, Hanna Hemeling, Phyllis Stöhr, Jürgen Beck, Andreas Vlachos

## Abstract

Homeostatic synaptic plasticity serves to maintain neuronal function within a dynamic range upon perturbations in network activity. While coordinated structural and functional changes at synaptic sites play a crucial role in adaptive processes, the specific regulatory mechanisms and biological relevance of homeostatic plasticity in the human brain warrant further investigation. In this study, we investigated the effects of neural network silencing, achieved through pharmacological inhibition of voltage-gated sodium channels or glutamatergic neurotransmission – common targets of antiepileptic medication – on functional and structural properties of murine and human cortical tissue. Using mouse entorhino-hippocampal tissue cultures, acute neocortical slices of adult mice, and human brain tissue, we characterize homeostatic synaptic plasticity across models, brain regions, and species. Our findings demonstrate local homeostatic synaptic plasticity in the adult human neocortex, highlighting the potential effects of antiepileptic medication in brain regions unaffected by the primary diseases, which might represent a mechanism for neuropsychiatric effects linked to these medications and increased seizure susceptibility upon discontinuation of antiepileptic medication.

## INTRODUCTION

Homeostatic synaptic plasticity plays a crucial role in maintaining the activity of neurons within a dynamic range (Davis and Bezprozvanny, 2001; Turrigiano, 2012). Its contribution to network stability and associative plasticity have been acknowledged in numerous computational and experimental studies (Li et al., 2019; Vercruysse et al., 2021; Wen and Turrigiano, 2021; Gallinaro et al., 2022). Although synaptic homeostasis has been extensively investigated in cell culture models and rodent brains, its presence in the adult human neocortex has yet to be demonstrated.

The use of tetrodotoxin (TTX), a voltage-gated sodium channel inhibitor that suppresses network activity, has emerged as a prevalent *in vitro* model for studying homeostatic synaptic plasticity (Turrigiano et al., 1998; Gainey et al., 2009; Ratkai et al., 2021). In addition to its effectiveness as a neurotoxin, TTX exerts potent effects on network properties. This raises the question about the extent to which structural, functional, and molecular adaptations identified from TTX-induced network silencing can be extrapolated to homeostatic plasticity occurring in the intact (human) brain under normal and pathological conditions.

In this study, we investigated the impact of antiepileptic medication on homeostatic synaptic plasticity. Our analysis encompassed human neocortical resections obtained from patients with tumors and epilepsy. Specifically, we examined whether the use of antiepileptic drugs, such as lamotrigine (LTG), in a patient’s medical history correlated with changes in excitatory synaptic strength of superficial pyramidal neurons. Our findings revealed that patients receiving antiepileptic medication exhibited enhanced excitatory synaptic transmission compared to untreated patients. Excitatory synaptic plasticity in the human neocortex was accompanied by structural changes in dendritic spines and modifications in the cortical transcriptome. The observed changes in the human neocortex were consistent with analyses conducted in acute slices prepared from the mouse medial prefrontal cortex (mPFC), as well as in mouse entorhino-hippocampal tissue cultures treated with LTG or TTX. These results demonstrate that antiepileptic drugs induce homeostatic synaptic plasticity in the adult human neocortex – which can be well reflected in both *in vitro* and *in vivo* mouse models – with significant implications concerning the neuropsychiatric effects linked to these medications and the increased seizure susceptibility upon discontinuing antiepileptic treatment.

## MATERIALS AND METHODS

### Ethics statement

Human tissue was received via a local biobank of the Department for Neurosurgery at the Faculty of Medicine, University of Freiburg (AZ 472/15_160880). All experiments were approved by the Local Ethics Committee, University of Freiburg (AZ 593/19). Patients gave their informed consent to use the neurosurgical resection material for research purposes.

Experimental procedures with animals were performed according to German animal welfare legislation and approved by the appropriate animal welfare committee and the animal welfare officer of the Albert-Ludwigs-University Freiburg, Faculty of Medicine (X17-07/K, X21-01/B; preparation of organotypic tissue cultures) or the competent authorities (G22-025; *in vivo* AED treatment). Mice were maintained in a 12 h light/dark cycle with food and water available *ad libitum*. Every effort was made to minimize distress and pain of animals.

### Preparation of tissue cultures

Organotypic entorhino-hippocampal tissue cultures were prepared from C57BL/6J mice of either sex at postnatal day 3-5 as previously described (Del Turco and Deller, 2007). The cultivation medium contained 50% (v/v) MEM, 25% (v/v) basal medium eagle, 25% (v/v) heat-inactivated normal horse serum, 25 mM HEPES buffer solution, 0.15% (w/v) bicarbonate, 0.65% (w/v) glucose, 0.1 mg/ml streptomycin, 100 U/ml penicillin, and 2 mM glutamax. The pH was adjusted to 7.3 and the medium was replaced 3 times per week. Prior to experimental procedures, all tissue cultures were allowed to mature in a humidified atmosphere with 5% CO_2_ at 35 °C for at least 18 days.

### Pharmacology

For some experiments, tissue cultures were treated with tetrodotoxin (TTX, 2 µM, 2 d; #ab120055, Abcam) while control cultures were only treated with vehicle (water, 1 µl). For experiments with antiepileptic drugs, tissue cultures were treated with lamotrigine (LTG, 100 µM, 2 d), and again control cultures were treated with vehicle (0.1% (v/v) DMSO, 2 d). For *in vivo* experiments, adult mice were randomly assigned to either the treatment or control group. Subsequently, they were injected intraperitoneally with LTG (20 mg/kg in 5% (v/v) DMSO in corn oil, 2 d) or vehicle-only (5% (v/v) DMSO in corn oil, 2 d). After injection, all mice acted normally and did not show any signs of pathological behavior.

### Preparation of acute mouse cortical slices

Adult mice of either sex (C57BL/6J; 7-weeks old, adapted to housing conditions) were treated for two days with LTG (20 mg/kg, 5% (v/v) DMSO in corn oil, daily intraperitoneal injections) and subsequently used for the preparation of acute cortical slices. After two days, animals were anesthetized with ketamine (100 mg/kg) / xylazine (20 mg/kg), rapidly decapitated and the brain was removed. For slicing, the brain was embedded in low-melting-point agarose (Sigma-Aldrich #A9517; 1.8% w/v in PBS) tilted dorsally at a 15° angle. Coronal sections (350 µm) containing the medial prefrontal cortex (mPFC) were prepared using a Leica VT1200S vibratome in chilled NMDG-aCSF containing (in mM) 92 NMDG, 2.5 KCl, 1.25 NaH_2_PO_4_, 30 NaHCO_3_, 20 HEPES, 25 glucose, 2 thiourea, 5 Na-ascorbate, 3 Na-pyruvate, 0.5 CaCl_2_, and 10 MgSO_4_, (pH = 7.3-7.4). Subsequently, slices were transferred to cell strainers with 40 µm pores in NMDG-aCSF at 34°C. Sodium levels were gradually increased as previously described and validated (Ting et al., 2018; Lenz et al., 2021b). After recovery, slices were maintained in an extracellular solution containing (in mM) 92 NaCl, 2.5 KCl, 1.25 NaH_2_PO_4_, 30 NaHCO_3_, 20 HEPES, 25 glucose, 2 thiourea, 5 Na-ascorbate, 3 Na-pyruvate, 2 CaCl_2_, and 2 MgSO_4_ (pH = 7.3-7.4) at room temperature until further experimental assessment. At all times, the used medium was oxygenated continuously (5% CO_2_/95% O_2_).

### Preparation of acute human cortical slices

Cortical resections for therapeutic or procedural indications, i.e. cortical access tissue, were performed as previously described (Straehle et al., 2023). Cortical tissue was transferred to chilled and carbo-oxygenated NMDG-aCSF (Lenz et al., 2021a). Subsequently, it was embedded in low-melting-point agarose and 400 µm sections were cut with a Leica VT1200S vibratome perpendicular to the pial surface in chilled NMDG-aCSF. Transfer, recovery, and maintenance of human slices were performed as described above for acute mouse cortical slices and in line with a previously published protocol (Ting et al., 2018). At all times, the used medium was oxygenated continuously (5% CO_2_/95% O_2_). None of the human cortical slices showed any macroscopically visible pathologies.

### Whole-cell patch-clamp recordings

Whole-cell patch-clamp recordings in organotypic tissue cultures were performed in a bath solution containing (in mM) 126 NaCl, 2.5 KCl, 26 NaHCO_3_, 1.25 NaH_2_PO_4_, 2 CaCl_2_, 2 MgCl_2_, and 10 glucose. In order to record miniature excitatory postsynaptic currents (mEPSCs), the solution was substituted with TTX (0.5 µM; #ab120055, Abcam), D-AP5 (10 µM; #ab120003, Abcam) and (−)-bicuculline-methiodide (10 µM; #ab120108, Abcam). For recordings of spontaneous EPSCs (sEPSCs) in acute murine and human cortical slices, the bath solution contained (in mM) 92 NaCl, 2.5 KCl, 1.25 NaH_2_PO_4_, 30 NaHCO_3_, 20 HEPES, 25 glucose, 2 thiourea, 5 Na-ascorbate, 3 Na-pyruvate, 2 CaCl_2_, and 2 MgSO_4_ (pH = 7.3-7.4). Regardless of the medium, recordings were carried out at 35° under continuous oxygenation (5% CO_2_/95% O_2_) and 3-6 cells were patched per culture or slice. Cells were visually identified using an LN-Scope (Luigs and Neumann, Germany) equipped with infrared dot-contrast and a 40× water-immersion objective (numerical aperture [NA] 0.8; Olympus). Human superficial (layer 2/3) pyramidal cells were visualized on the pia-white matter axis at a distance of 500-1000 µm to the pial surface. Electrophysiological signals were amplified using a Multiclamp 700B amplifier, digitized with a Digidata 1550B digitizer, and visualized with the pClamp 11 software package. Patch pipettes contained (in mM) 126 K-gluconate, 4 KCl, 10 HEPES, 4 MgATP, 0.3 Na_2_GTP, 10 PO-creatine, and 0.3% (w/v) biocytin (pH = 7.25 with KOH; 285 mOsm/kg) and had a tip resistance of 3-5 MΩ. For both mEPSC and sEPSC recordings, cells were recorded in voltage-clamp mode at a holding potential of −70 mV. Series resistance was monitored before and after recording and recordings were discarded if the resistance reached ≥ 30 MΩ. In cortical neurons of acute murine and human slices, intrinsic cellular properties were recorded in current-clamp mode following sEPSC recordings. Pipette capacitance of 2.0 pF was corrected and series resistance was compensated using the automated bridge balance tool of the MultiClamp commander. I-V-curves were generated by injecting 1 s square pulse currents starting at - 100 pA and increasing in 10 pA steps until +500 pA current injection was reached with a sweep duration of 2 s. For human pyramidal neurons, the current was increased up to +1100 pA. Again, series resistance was monitored and recordings were discarded if the resistance reached ≥ 30 MΩ.

### Immunofluorescence and post-hoc staining

After electrophysiological assessment, tissue cultures or cortical slices were fixed in 4% (w/v) paraformaldehyde (PFA; in PBS (0.1 M, pH 7.4) with 4% (w/v) sucrose) overnight. After fixation, they were washed in PBS (0.1 M, pH 7.4) and incubated with 10% (v/v) normal goat serum (NGS) in 0.5% (v/v) Triton X-100 containing PBS for 1 h, in order to reduce nonspecific staining. For immunofluorescence, slices were incubated with an anti-NeuN antibody (Rabbit polyclonal, 1:1000; #ab104225, Abcam) overnight in 10% (v/v) normal goat serum (NGS) in 0.1% (v/v) Triton X-100 containing PBS at 4° C overnight. After washing, suitable secondary antibodies (goat anti rabbit, Alexa Fluor Plus 555, #A-32732, Invitrogen) were added during the post-hoc visualization of patched neurons, for which the tissue was incubated with Streptavidin Alexa Fluor 488 (Streptavidin A488, 1:1000; #S32354 Invitrogen) diluted in 10% (v/v) normal goat serum (NGS) in 0.1% (v/v) Triton X-100 containing PBS at 4° C overnight. After washing, the tissue was incubated with DAPI nuclear stain (1:5000 in PBS for 15 minutes; #62248 Thermo Scientific) in order to visualize cytoarchitecture, transferred onto glass slides and mounted with a fluorescence anti-fading mounting medium (DAKO Fluoromount; #S302380-2, Agilent). In order to reduce age-related autofluorescence in human cortical slices, they were additionally incubated with Sudan black (0.1% [w/v] in 70% ethanol) for 10 min before the DAPI staining.

### Confocal imaging

For visualization of patched human cortical neurons, confocal images were acquired using a Leica SP8 laser-scanning microscope equipped with a 20× multi-immersion (NA 0.75; Leica), a 40× oil-immersion (NA 1.30; Leica), and a 63× oil-immersion objective (NA 1.40; Leica). For structural analysis of human cortical spines, an overview image of each cell was generated with the 40× oil-immersion objective setting the limits of the z-axis approximately 10 µm in each direction from the middle of the soma (z-step: 0.6 µm). Apical and basal dendritic segments were randomly chosen and their position tagged in the overview image, in order to be able to analyze their distance to soma. 5-25 segments of each cell were recorded using the 63× oil-immersion objective (resolution: 2048 x 2048 pixels, zoom: 4x, speed: 200 Hz, pinhole: 0.6 AU, z-step: 0.1 µm). Laser intensity and detector gain were set individually, in order to achieve comparable overall fluorescence intensities throughout stacks. Confocal image stacks were stored as .tif files.

### Transcriptome analysis

For experiments in organotypic tissue cultures, 5-6 cultures of the same mouse were collected and used per sample. For *in vivo* experiments, the mPFC of treated and untreated adult mice was dissected and the mPFC of the left hemisphere was used as one sample. In experiments with human material, the tissue of each patient was seen as one biological sample. RNA was isolated using the Monarch^®^ Total RNA Miniprep Kit (#T2010S; New England Biolabs) according to the manufacturer’s instructions. RNA quantity and quality were assessed using an Agilent RNA 6000 Pico Kit (#5067-1513; Agilent) with a 2100 Bioanalyzer (#G2939BA; Agilent). After RNA isolation from TTX-treated tissue cultures, library preparation and RNA sequencing was performed using the genome sequencer Illumina HiSeq technology in NovaSeq 6000 S4 PE150 XP sequencing mode (service provided by Eurofins). For further analysis, .fastq files were provided. All samples contained more than 10 M high-quality reads in total having at least a phred quality of 30 (>90% of total reads). After RNA isolation from the mPFC of mouse *in vivo* experiments and human neocortical samples, poly(A)-selection was performed using a poly(A)-selection mRNA magnetic isolation module (#E7490; New England Biolabs) according to the manufacturer’s instructions. For non-directional library preparation, NEBNext Ultra II RNA Library Preparation Kit for Illumina (#E7770; New England Biolabs) was used. Libraries were finally cleaned up with 0.8X SPRI beads following a standard bead purification protocol. Library purity and size distribution were assessed with a High Sensitivity DNA assay on a Bioanalyzer instrument (Agilent). The libraries were quantified using the NEBNext Library Quant Kit for Illumina (#E7630; New England Biolabs) based on the mean insert size provided by the Bioanalyzer. A 10 nM sequencing pool (120 μl in Tris-HCl, pH 8.5) was generated for sequencing on the NovaSeq6000 sequencing platform (Illumina; service provided by CeGaT GmbH, Tübingen, Germany). Paired-end sequencing with 150 bp read length was performed. Data were analyzed at the Galaxy platform (usegalaxy.eu; (Galaxy, 2022)). All files contained more than 10 M high-quality reads (after mapping to the reference genome) with a phred quality of at least 30 (>90% of total reads).

### Quantification and statistics

Electrophysiological data were assessed using the pClamp 11 software package (Axon Instruments). mEPSC and sEPSC properties were analyzed with an automated templated-based search tool for event detection. The template was previously created in the Clampfit11 software based on manually identified and selected EPSCs. The template match threshold was set at 2.5 and the time period for the allowed duration of a single event starting from the baseline was defined. Both parameters were not changed within the different recordings or groups. For recordings of pyramidal neurons of acute murine and human slices, the resting membrane potential was assessed from the baseline value of the I-V-curve. Input resistance was calculated for the injection of −100 pA current at a time frame of 200 ms with a maximum distance to the initial hyperpolarization.

Structural analysis of spine density and volume of human pyramidal neurons was conducted by investigators blinded to experimental conditions. Spine density was assessed by counting spines manually in the z-stack of the confocal images with the ImageJ software package (https://imagej.nih.gov/ij/) and normalizing the number to the segment’s length. The distance to soma was measured based on the segment’s marked position in the respective cell’s overview image. Spine volume was analyzed using the surface detection tool of the Imaris 9.5 software package (Oxford Instruments).

For transcriptome analysis, sequencing data were uploaded to the Galaxy web platform (public server: usegalaxy.eu), and the analysis was performed based on the RNA-seq data analysis tutorial (Batut et al., 2021). In short, the CUTADAPT tool was used to remove adapter sequencing, low quality and short reads. Remaining reads were subsequently mapped using the RNA STAR tool with the mm10 (Mus musculus) or the hg38 (human) reference genome. The evidence-based annotation of the mouse genome (GRCm38), version M25 (Ensembl 100), or the human genome (GRCh38), version 40 (Ensembl 106), served as a gene model (GENCODE). An unstranded FEATURECOUNT analysis of the RNA STAR output was performed for an initial assessment of gene expression. Only samples that contained > 60% uniquely mapping reads (feature: “exon”) were considered for further analysis. Statistical evaluation was performed using DESeq2 with “treatment” as the primary factor affecting gene expression. Genes with mean reads (base mean) < 150 were excluded from further analysis. Genes with an adjusted p-value of < 0.05 were considered as differentially expressed. For data visualization, a modified version of a previously published workflow was used (Reimand et al., 2019). The g:Profiler (version e107_eg54_p17_bf42210) with g:SCS multiple testing correction method (significance threshold of 0.05 (Raudvere et al., 2019)) was used for functional enrichment analysis. Heatmaps were generated based on the z-scores of the normalized count tables.

Data were statistically analyzed using GraphPad Prism 9 (GraphPad Software, USA). All values represent mean ± standard error of the mean (s.e.m.). In the graphs demonstrating spine volumes, the error bars depict the 5.-95. percentile while the data points show the results outside that range. In these experiments, a Kruskal-Wallis test was used for statistical comparisons. Otherwise, a non-parametric Mann-Whitney test was employed for comparison of two experimental groups. XY-plots were statistically assessed using a repeated measure (RM) two-way ANOVA test with Sidak’s (two groups) multiple comparisons. P-values < 0.05 were considered statistically significant (*p < 0.05, **p < 0.01, ***p < 0.001); results without statistical significance were indicated as ‘ns’. Statistical differences in XY-plots were indicated in the legend of the figure panels (*), irrespective of their localization and the level of significance. N-numbers are provided in the figure legends.

### Data availability

Sequencing data have been deposited in the Gene Expression Omnibus (GEO) repository (accession number: GSE244095). Original data are available upon reasonable request.

### Digital illustrations

Confocal images were stored as .tif files and image brightness and contrast were adjusted. Figures were prepared using the ImageJ software package (https://imagej.nih.gov/ij/) and Photoshop graphics software (Adobe, San Jose, CA, USA).

## RESULTS

### Voltage-gated sodium channel inhibition induces plasticity in organotypic entorhino-hippocampal tissue cultures

Mouse organotypic entorhino-hippocampal tissue cultures were treated for two days with 2 µM TTX (Figure 1A). A significant advantage of these tissue cultures is their laminar organization, enabling different cell types, including dentate granule cells (dGC) and CA1 pyramidal neurons (CA1-PC), to receive innervation through the perforant path and intra-hippocampal pathways (Figure 1B). In order to characterize the effects of TTX-induced inhibition of voltage-gated sodium channels, we conducted a transcriptome analysis of whole tissue cultures (Figure 1C). A substantial number of genes exhibited differential expression in response to TTX treatment compared to control cultures (UP = 2182 genes, DOWN = 2240 genes). The gene set enrichment analysis (Figure S1, g:Profiler analysis) focused on gene ontology categories such as cellular compartment (GOCC) and biological process (GOBP), revealed differential expression of synapse and plasticity-related gene sets, including presynaptic, postsynaptic and synaptic vesicle-related genes (Figure 1D).

**Figure 1:**
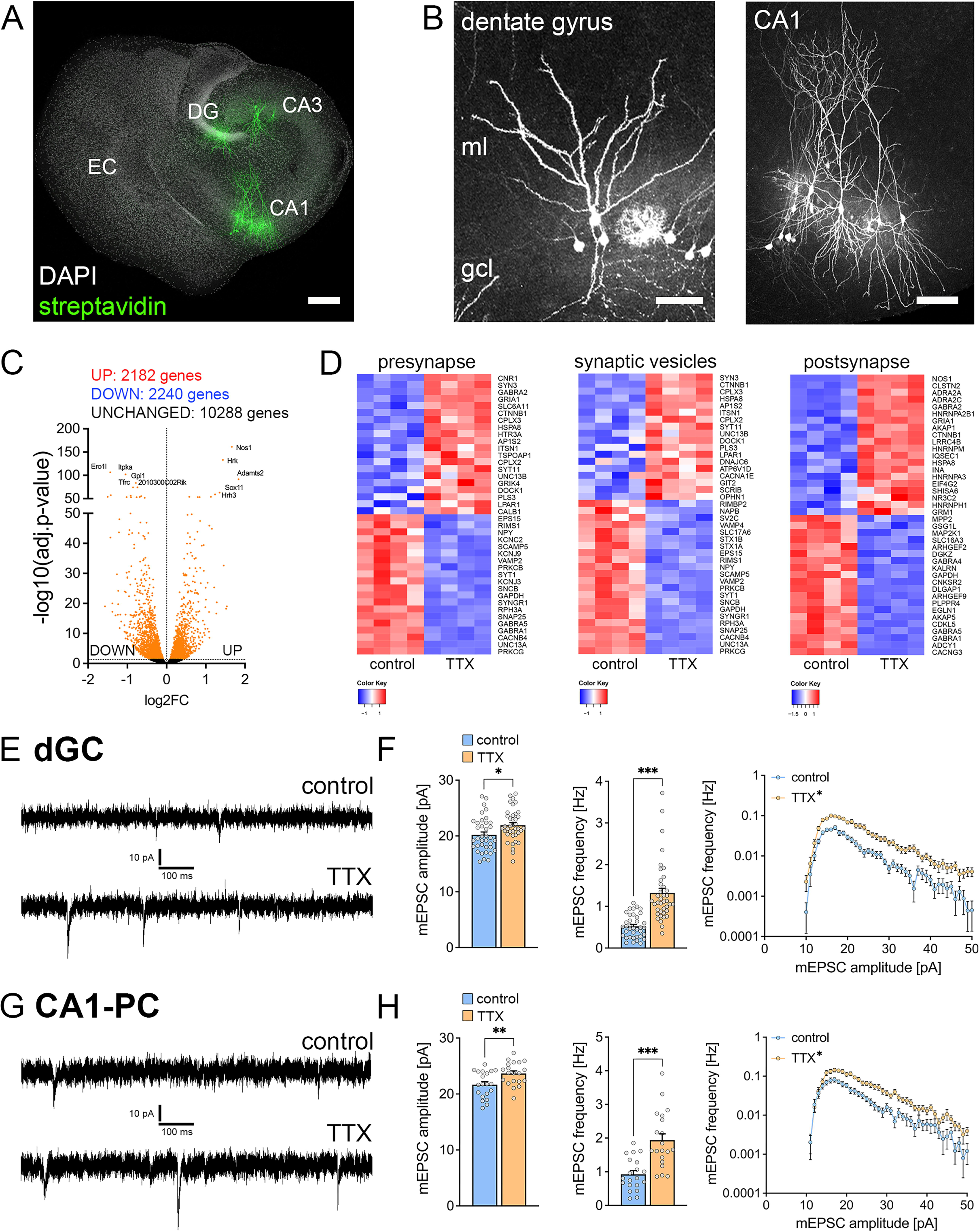
TTX-treatment induces synaptic plasticity in organotypic tissue cultures. (A) Sample image of an organotypic entorhino-hippocampal tissue culture stained with DAPI nuclear stain and post-hoc visualization of dentate granule cells and pyramidal neurons in CA1 and CA3. EC, entorhinal cortex; DG, dentate gyrus. Scale bar, 250µm. (B) Post-hoc stained dentate granule cells (left image) and CA1 pyramidal neurons (right image) in organotypic entorhino-hippocampal tissue cultures. gcl, granule cell layer; ml, molecular layer; Scale bar dentate gyrus, 50 µm; Scale bar CA1, 100 µm. (C)-(D) RNA-Sequencing and transcriptome analysis of control cultures and cultures treated with TTX (2 µM, 2 days). (C) The volcano plot represents fold change and adjusted p-value of differentially expressed genes (indicated in orange, the Top-10 differentially expressed genes are labeled). (D) Heat maps of differentially expressed genes assigned to the GOCC terms “presynapse” (left panel), “synaptic vesicles” (panel in the middle), and “postsynapse” (right panel). (E)-(F) Sample traces and group data of whole-cell patch-clamp recordings of mEPSCs in dentate granule cells (dGC). Treatment with TTX (2 µM, 2 days) induces synaptic strengthening reflected by a significant increase in mEPSC amplitude and frequency (F, n_control_ = 37 cells in 7 cultures, n_TTX_ = 39 cells in 7 cultures; Mann-Whitney test, RM two-way ANOVA for amplitude/frequency analysis). (G)-(H) Sample traces and group data of whole-cell patch-clamp recordings of mEPSCs in CA1 pyramidal cells (CA1-PC). Treatment with TTX (2 µM, 2 days) induces synaptic strengthening reflected by a significant increase in mEPSC amplitude and frequency (F, n = 20 cells in 4 cultures in each condition respectively; Mann-Whitney test, RM two-way ANOVA for amplitude/frequency analysis). Individual data points are indicated by gray dots. Values represent mean ± s.e.m. (*p < 0.05; **p < 0.01; ***p < 0.001).

Furthermore, we assessed the functional effects of TTX treatment by recording α-Amino-3-hydroxy-5-methylisoxazol-4-propionsäure (AMPA) receptor mediated mEPSCs in both dGCs and CA1-PCs (Figure 1E, G). TTX treatment resulted in a significant increase in mEPSC amplitudes and frequencies in the dGCs (Figure 1F). Similarly, TTX treatment led to an increase in mean mEPSC amplitude and frequency in CA1-PCs (Figure 1H). Notably, excitatory synaptic strengthening in CA1-PCs was observed only when bicuculline-methiodide was used to block network inhibition during the recordings (Figure S2). We conclude that TTX-mediated inhibition of voltage-gated sodium channels induces excitatory synaptic plasticity in different cell types, namely GCs and CA1-PCs, within organotypic entorhino-hippcampal tissue cultures.

### Lamotrigine (LTG) induces strengthening of excitatory neurotransmission in organotypic entorhino-hippocampal tissue cultures

As partial voltage-gated sodium channel inhibition is a well-established mechanism of antiepileptic medication for treating primary or secondary epileptic seizures, we assessed the potential of the commonly used antiepileptic drug LTG to induce a compensatory synaptic strengthening in mouse organotypic entorhino-hippocampal tissue cultures (Figure 2A-D). After a 2-day treatment with LTG (100 µM) or vehicle-only, we recorded AMPA receptor-mediated mEPSCs in both dGC (Figure 2A, B) and CA1-PCs (Figure 2C, D). LTG-treatment induced changes in excitatory synaptic transmission in both cell types. In dGCs, a significant increase in mEPSC frequencies was detected, while mEPSC amplitudes remained unchanged (Figure 2A). Additionally, the amplitude/frequency-plot revealed a significant excitatory synaptic strengthening upon LTG treatment (Figure 2B). In contrast, CA1-PCs exhibited a significant increase in mEPSC amplitudes (Figure 2C), with no differences observed in mEPSC frequencies or the amplitude/frequency-plot (Figure 2D). Although these findings differ from the effects of complete network inhibition by TTX, we concluded that a partial voltage-gated sodium channel inhibition induced by LTG promotes excitatory synaptic plasticity in organotypic entorhino-hippocampal tissue cultures.

**Figure 2:**
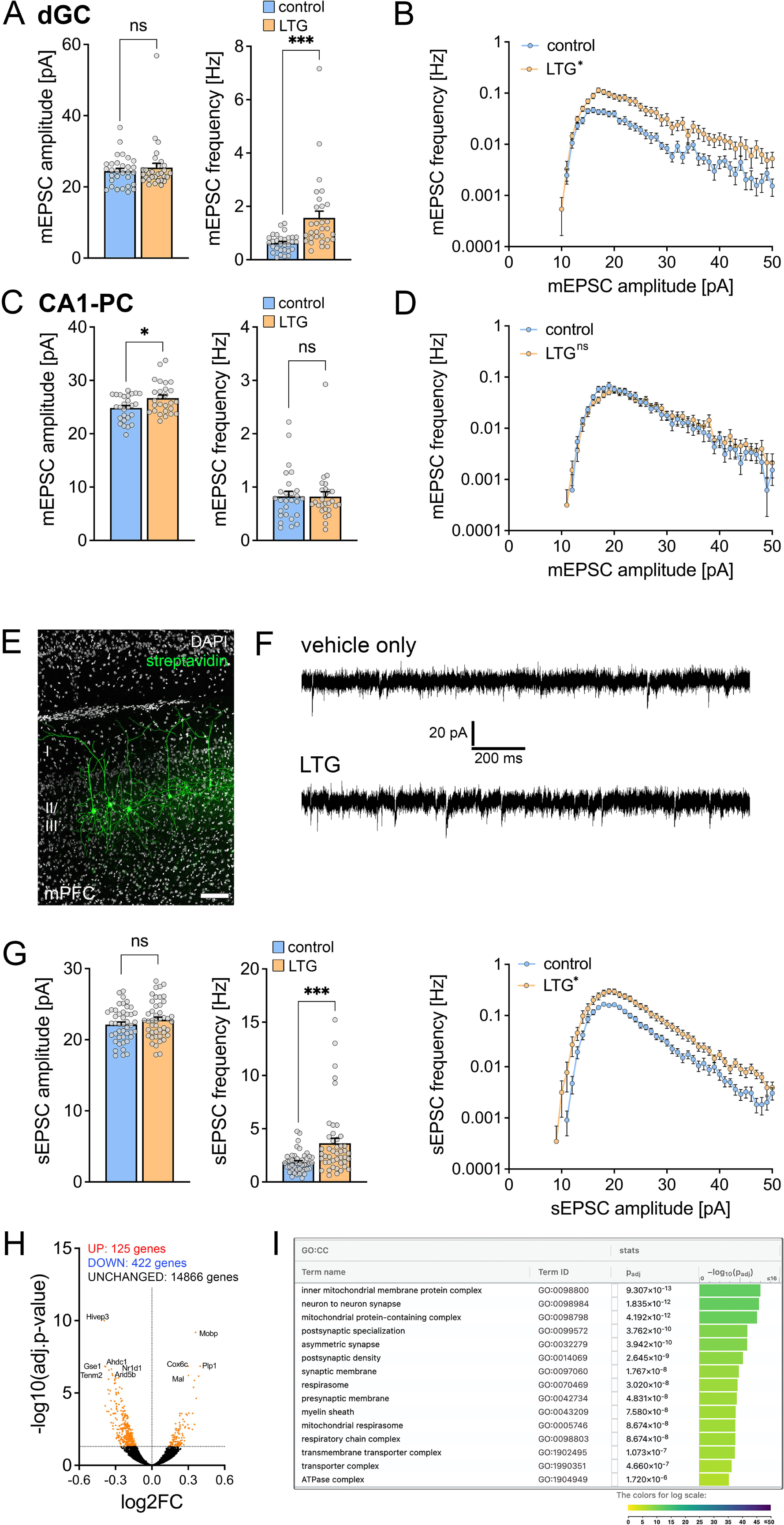
Lamotrigine induces synaptic strengthening in organotypic tissue cultures and the mPFC of adult mice. (A)-(B) Group data of whole-cell patch-clamp recordings of mEPSCs in dGCs in control cultures and cultures treated with Lamotrigine (100 µM, 2 days). Treatment with Lamotrigine induces a significant increase in mEPSC frequency (A, n = 30 cells in 5 cultures in each condition respectively; Mann-Whitney test, RM two-way ANOVA for amplitude/frequency analysis). (C)-(D) Group data of whole-cell patch-clamp recordings of mEPSCs in CA1-PCs in control cultures and cultures treated with Lamotrigine (100 µM, 2 days). Treatment with Lamotrigine induces a significant increase in mEPSC amplitude (C, n = 27 cells in 5 cultures for each condition respectively; Mann-Whitney test, RM two-way ANOVA for amplitude/frequency analysis). (E) Example image of post-hoc stained pyramidal neurons in layer 2/3 of the mPFC of ∼7 week old mice. Scale bar, 100 µm. (F, G) Sample traces (F) and group data (G) of whole-cell patch-clamp recordings of sEPSCs in layer 2/3 pyramidal neurons of control mice and mice treated with Lamotrigine (20 mg/kg in 5% (v/v) DMSO in corn oil, 2 days). Treatment with Lamotrigine induces homeostatic plasticity reflected by a significant increase of the sEPSCs frequency (n = 45 cells in 4 mice for each condition respectively; Mann-Whitney test, RM two-way ANOVA for amplitude/frequency analysis). (H)-(I) RNA-Sequencing and transcriptome analysis of isolated mPFC from ∼7 week old control mice and mice treated with lamotrigine (20 mg/kg in 5% (v/v) DMSO in corn oil, 2 days). (H) Volcano plot represents fold change and adjusted p-value of differentially expressed genes (indicated in orange, the Top-10 differentially expressed genes are labeled; n_control_ = 4 mice, n_LTG_ = 3 mice). (I) Top-10 Gene Ontology Cellular Component (GOCC) regulated terms in the mPFC. The color code represents the - log10(p-adj) of the differentially expressed genes. Individual data points are indicated by gray dots. Values represent mean ± s.e.m. (*p < 0.05; ***p < 0.001; ns, non-significant difference).

### *In vivo* LTG treatment elicits excitatory synaptic strengthening in superficial pyramidal neurons of the medial prefrontal cortex in adult mice

In order to investigate homeostatic synaptic plasticity through LTG in the neocortex of adult mice, we administered LTG (100 mg/kg, i.p. daily injection for 2 days) or vehicle-only (5% (w/v) DMSO in corn oil) to adult C57BL/6J mice of both sexes. Subsequently, we assessed excitatory neurotransmission in superficial (layer 2/3) pyramidal neurons of the (mPFC) by recording AMPA-receptor mediated sEPSCs (Figure 2E, F). After 2 days of treatment, we observed a significant increase in sEPSC amplitudes, whereas sEPSC frequencies remained unchanged (Figure 2G). The amplitude/frequency-plot confirmed excitatory synaptic strengthening following *in vivo* LTG treatment (Figure 2G).

To correlate functional changes in excitatory neurotransmission with gene expression (Figure 2H, I), transcriptome analysis of the mPFC was conducted. Differential gene expression was observed between LTG-treated and vehicle-only groups (Figure 2H; UP = 125 genes, DOWN = 422 genes). Notably, gene set enrichment analysis revealed an enrichment of both synaptic and mitochondrial gene sets (Figure 2I). Although bulk-RNASeq data sets have limitations, these findings identify transcriptome changes corresponding to the functional changes in excitatory neurotransmission in the mouse mPFC of LTG-treated adult mice. We conclude that LTG induces excitatory synaptic plasticity in the mPFC of adult mice.

### Antiepileptic treatment induces homeostatic synaptic plasticity in the adult human neocortex

Building upon our *in vitro* and *in vivo* LTG experiments, we investigated whether patients receiving antiepileptic medication displayed enhanced excitatory neurotransmission in neocortical regions. Using neurosurgical access tissue from tumor or epilepsy patients (Table S1), we prepared acute neocortical slices (Figure 3). Due to individual clinical conditions, a variety of antiepileptic drugs was found in the medical records of patients, such as lamotrigine, lacosamide, brivaracetam, levetiracetam, perampanel, or no antiepileptic medication was found in their medical records. However, all antiepileptic drugs aim at dampening network activity to achieve seizure control. AMPA-receptor mediated sEPSCs were recorded from layer 2/3 pyramidal neurons (Figure 3A, B). Comparative analysis of acute slices from patients receiving antiepileptic therapy versus those without revealed increased sEPSC amplitudes, half widths, and frequencies in layer 2/3 pyramidal neurons (Figure 3C). The amplitude/frequency-plot further confirmed an increased occurrence of sEPSC events with higher amplitudes (Figure 3C). Thus, we conclude that antiepileptic treatment strengthens excitatory neurotransmission in the adult human neocortex, providing evidence for homeostatic synaptic plasticity at the single-cell level in the human brain.

**Figure 3:**
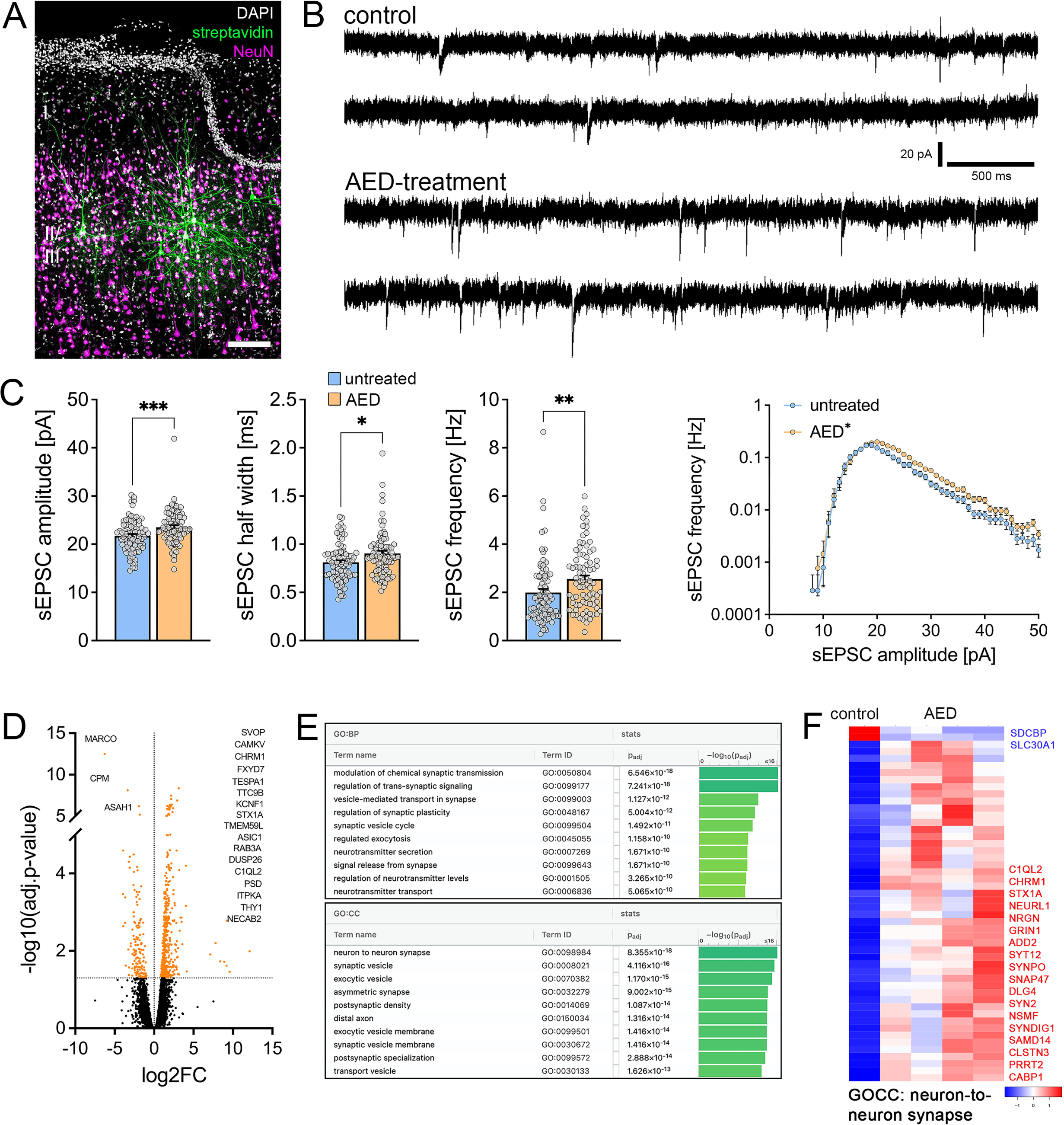
Treatment with antiepileptic drugs (AED) is associated with synaptic strengthening in the human cortex. (A) Example image of post-hoc stained superficial pyramidal neurons in the human cortex (green). Neuronal cell bodies are visualized by a staining against NeuN (magenta). Scale bar, 150 µm. (B)-(C) Sample traces (B) and group data of sEPSCs recorded from superficial pyramidal neurons in the cortical samples of control patients and patients treated with antiepileptic drugs (AED, C, D). Recordings in the cortex of AED-treated patients show a significantly higher sEPSC amplitude, half with and frequency (n_control_ = 81 cells in 3 samples, n_AED_ = 80 cells in 4 samples; Mann-Whitney test, RM two-way ANOVA for amplitude/frequency analysis). (D)-(F) RNA-sequencing of cortical tissue of control patients and patients treated with antiepileptic drugs (n_control_ = 1 sample, n_AED_ = 4 samples). (D) The volcano plot represents fold change and adjusted p-value of differentially expressed genes (indicated in orange, the Top-20 differentially expressed genes are named). (E) Top-10 Gene-Ontology Cellular Component (GOCC) and Gene-Ontology Biological Process (GOBP) regulated terms. The color code represents the -log10(p-adj) of the enrichment. (F) Heatmap based on z-scores of normalized counts from differentially expressed genes of GOCC: neuron- to-neuron synapse; subset of genes indicated on the side. Individual data points are indicated by gray dots. Values represent mean ± s.e.m. (*p < 0.05; **p < 0.01; ***p < 0.001).

### Antiepileptic treatment induces transcriptome changes in the neocortex of human patients

We conducted a transcriptome analysis to gain deeper insights regarding the effects of antiepileptic treatment on the human neocortex. By comparing cortical samples from neurosurgical access material of patients with antiepileptic treatment and without, we identified a significant number of genes with differential expression (Figure 3D; UP = 497 genes, DOWN = 154 genes, UNCHANGED = 11323 genes). Gene set enrichment analysis highlighted an enrichment of synaptic/neuronal differential gene expression (Figure 3E), such as ‘modulation of chemical synaptic transmission’, ‘neuron to neuron synapse’, and ‘synaptic vesicle’ (Figure 3F; gene ontology: cellular compartment, GOCC for neuron-to-neuron synapse). These findings suggest that antiepileptic treatment is associated with coordinated functional and transcriptomic changes in the human neocortex.

### Antiepileptic drug treatment increases spine head volumes at distinct locations along the dendritic tree of superficial pyramidal neurons in the human neocortex

We examined the potential impact of antiepileptic drug treatment on the structural plasticity of dendritic spines in human superficial pyramidal neurons, which were *post hoc* visualized after the assessment of excitatory neurotransmission (Figure 4A). Dendritic spines were visualized in both apical and basal dendritic regions (Figure 4B), and we analyzed spine density and spine head volume. Initially, we assessed whether age influenced the structural features of dendritic spines (Figure S3). We observed a decline in dendritic spine density during aging in both apical and basal dendrites, while spine head volume remained constant (Figure S3). To eliminate “age” as a potential confounding factor, we focused on investigating the effects of antiepileptic treatment specifically on dendritic spine head volume.

**Figure 4:**
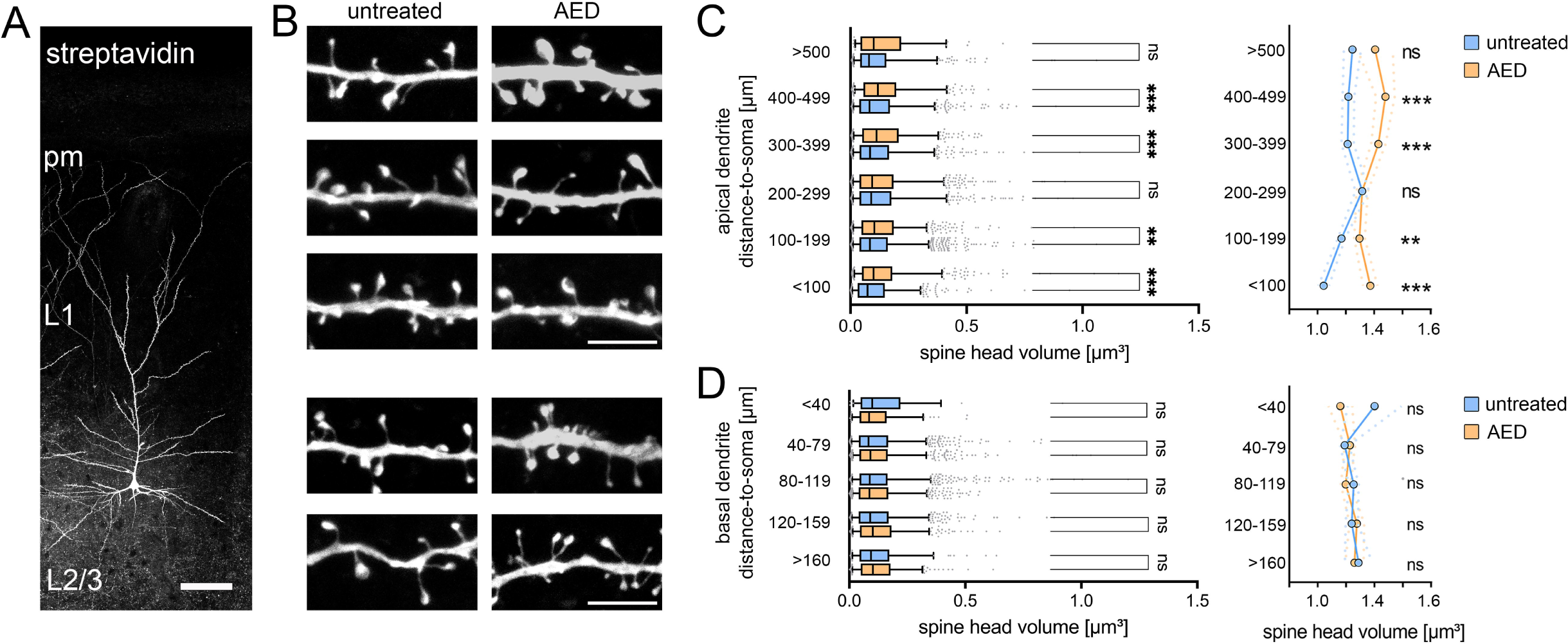
Treatment with antiepileptic drugs (AED) is associated with plastic changes of spine morphology in the human cortex. (A) Example image of a post-hoc stained superficial pyramidal neurons in the human cortex. L1, layer 1; L2/3, layer 2/3; pm, pia mater. Scale bar, 100 µm. (B) Example images of post-hoc stained apical and basal dendritic segments with spines of superficial pyramidal neurons in the cortex of patients without AED-treatment (untreated, images on the left) and patients with AED-treatment (AED, images on the right). (C) Spine volume of spines at the apical dendrites of superficial pyramidal neurons is increased at certain distances to the soma in samples of patients with AED-treatment (control, n_<100_ = 698, n_100-199_ = 1861, n_200-299_ = 785, n_300-399_ = 758, n_400-499_ = 847, n_>500_ = 464 spines in 3 samples; AED-treatment, n_<100_ = 668, n_100-199_ = 890, n_200-299_ = 1018, n_300-399_ = 518, n_400-499_ = 341, n_>500_ = 101 spines in 4 samples; Kruskal-Wallis test). (D) Spine volume of spines at the basal dendrites of superficial pyramidal neurons shows no difference between samples of patients without AED-treatment and patients with AED-treatment (control, n_<40_ = 59, n_40-79_ = 1152, n_80-119_ = 1367, n_120-159_ = 865, n_>160_ = 200 spines in 3 samples; AED treatment, n_<40_ = 135, n_40-79_ = 974, n_80-119_ = 927, n_120-159_ = 419, n_>160_ = 298 spines in 4 samples; Kruskal-Wallis test). Whiskers depict the 5.-95. percentile, individual dots demonstrate data points outside of that range (**p < 0.01; ***p < 0.001; ns, non-significant difference).

In apical dendrites, we observed an antiepileptic drug-related increase in spine head volumes that was dependent on the distance from the soma: Spines in close proximity and at longer distances from the soma showed increased spine head volumes, while spines at intermediate distances remained unaffected by antiepileptic drug (AED) treatment (Figure 4C). However, AED treatment did not induce any changes in dendritic spine head volumes of basal dendrites (Figure 4D). These findings demonstrate a compartment-specific structural synaptic adjustment in the adult human cortex, as AED-induced changes in excitatory synaptic strength involve coordinated structural and functional changes at distinct locations along the dendritic tree.

## DISCUSSION

The significant finding of this study lies in demonstrating excitatory synaptic adjustments within the adult human neocortex that is in line with the concept of homeostatic synaptic plasticity. Through analysis of acute slices from neurosurgical resections, we show that antiepileptic medication induced a compensatory strengthening of excitatory neurotransmission onto superficial pyramidal neurons of the neocortex. These functional changes align with transcriptome modifications and structural adjustments in dendritic spines, affecting apical dendrites while sparing basal dendrites. As a result, this study provides evidence for local homeostatic synaptic adjustment within the human neocortex. Our experiments corroborate previous findings, confirming that the inhibition of voltage-gated sodium channels triggers homeostatic synaptic plasticity across various brain regions (Turrigiano et al., 1998; Sutton et al., 2006; Dubes et al., 2022; Kavalali and Monteggia, 2023). Importantly, this plasticity arises within unaffected brain regions of healthy mouse brain tissue and in human brain tissue samples from patients, which received antiepileptic medication. These findings open novel avenues for enhancing our understanding of the biological significance of homeostatic synaptic plasticity in the human brain, spanning both healthy and diseased conditions. Initially characterized as a negative-feedback mechanisms that regulates all synapses of a neuron (referred to as synaptic scaling; (Turrigiano et al., 1998)), substantial evidence supports the notion that homeostatic synaptic plasticity can occur locally in specific sets of synapses (Kim and Tsien, 2008; Vlachos et al., 2013; Lenz et al., 2019). It is well established that pathological brain states, marked by modified network activity and denervation, induce homeostatic plasticity (Keck et al., 2013; Vlachos et al., 2013; Barnes et al., 2015; Lenz et al., 2023a). Although the biological importance of homeostatic plasticity remains a subject of discussion, its role in memory acquisition and specificity has recently been recognized (Wu et al., 2021). Furthermore, computational modeling indicates that homeostatic structural adaptations could potentially confer associative properties (Gallinaro et al., 2022; Lu et al., 2022). Still, a significant and unresolved matter within this field revolves around the intricate interplay between activity-dependent hebbian and homeostatic synaptic plasticity, i.e., positive and negative feed-back mechanism (Keck et al., 2013; Yee et al., 2017; Wu et al., 2022). These two processes share common downstream molecular mechanisms that recruit changes or adaptations in dendritic spine morphologies and synaptic neurotransmission, such as the synthesis, maturation, and trafficking of AMPA receptors (Turrigiano, 2008).

We examined dendritic spine morphologies to differentiate between global and local synaptic adjustments in somatic whole-cell recordings. Recent studies suggest that the morphological features of human dendritic spines are associated with distinct signal integration properties (Rosado et al., 2022; Hunt et al., 2023). Our study showed a contrast between basal and apical dendrites. Interestingly, apical dendrites exhibited increased spine volumes within the cohort of patients treated with AEDs, while basal dendrites remained unchanged. Hence, we infer that the phenomenon of homeostatic synaptic adjustment upon AED treatment is expressed at distinct synapses. Interestingly, even within the apical dendritic compartment not all dendrites showed significant differences between the groups. This observation offers the opportunity to investigate the capacity of distal apical, proximal apical and basal dendrites to manifest long-term potentiation and depression, and hence the metaplastic effects of homeostatic plasticity. The results are consistent with the emerging concept that dendrites within the human cortex could potentially carry out computations traditionally attributed to complex multilayered networks (Gidon et al., 2020).

Further investigation is required to identify the mechanisms driving AED-mediated local homeostatic synaptic plasticity. In a previous study, we explored the impact of all-trans retinoic acid (atRA) on human cortical tissue (Lenz et al., 2021a). atRA, a derivative of vitamin A, is associated with promoting homeostatic and Hebbian synaptic plasticity (Maghsoodi et al., 2008; Chen et al., 2014; Arendt et al., 2015; Li et al., 2019; Thapliyal et al., 2022). In this previous study, we observed increased excitatory synaptic strength and similar morphological adjustments in dendritic spines in acute human cortical slices under atRA exposure. These changes were sensitive to the pharmacological inhibition of mRNA translation but not DNA transcription ((Lenz et al., 2021a); c.f., (Sutton et al., 2006; Schanzenbacher et al., 2016)). It is pivotal to investigate whether the transcriptomic changes in this study connect with the depicted functional and structural homeostatic shifts. For instance, it is worth exploring whether the selective transport and stabilization of mRNAs into specific dendritic regions, particularly apical rather than basal, supports the outlined plastic changes. In any scenario, our experiments validate the occurrence of coordinated structural and functional adjustments in homeostasis within the intact human brain (c.f. (Althaus and Sutton, 2021)).

Human neocortical resections offer a distinctive pathway to explore neural characteristics within the human brain and to juxtapose these outcomes with observations obtained from animal models. Although the human brain shares similarities with the rodent brain (Verhoog et al., 2016; Lenz et al., 2021b), notable variations in signal integration and plasticity induction have been documented (Eyal et al., 2016; Gidon et al., 2020; Larkum et al., 2022). Our results demonstrate the manifestation of homeostatic synaptic plasticity in the human brain, underscoring its evolutionary preservation and significance across species, brain regions and research models. However, the diversity evident in medical records across human populations introduces potential confounding factors, such as additional medication. As revealed in this study, the aging process influenced spine density among superficial pyramidal neurons, while spine head volume remained stable. It is conceivable that other unidentified factors might have an additional impact on the results obtained from the neurosurgical material. Hence, a reverse translation approach was used to validate the effects of antiepileptic medication on cortical circuits within controlled *in vivo* and *in vitro* settings. Indeed, induction of homeostatic synaptic plasticity was observed in acute slices prepared from the mPFC of LTG treated mice and in entorhino-hippocampal slice cultures. These results confirm the utility of organotypic tissue cultures, which demonstrate *in vivo*-like layering (Maus et al., 2020; Lenz et al., 2023b), for deciphering mechanisms of synaptic plasticity, which are relevant in human neocortical circuits. Future systematic assessments across regions and species could unveil novel pharmacological targets to fine-tune homeostatic plasticity therapeutically.

Antiepileptic medication is commonly administered to manage seizures in tumor or epilepsy patients (Lattanzi et al., 2019; Moosa, 2019; Park et al., 2019; Verrotti et al., 2020; van der Meer et al., 2021). These medications target neurotransmitter receptors or voltage-gated channels, persistently reducing neural network activity. The immediate impact on neural networks can lead to pharmacologically-induced effects, like neuropsychiatric changes (Chen et al., 2016), triggering compensatory adaptations in neural networks. The extent to which these symptoms are causally linked to the kinetics of homeostatic synaptic plasticity induction, requires additional exploration. Of note, not all patients experience such effects. Given the regular administration of antiepileptic therapy, it is plausible that homeostatic synaptic strengthening persists throughout treatment. Importantly, abrupt discontinuation of medication often results in recurrent and exacerbated seizures (Wang et al., 2019a; Wang et al., 2019b). In this context, the excitatory synaptic plasticity induced by AEDs might play a crucial role: Elevated network activity following drug cessation could leverage heightened excitatory neurotransmission, potentially triggering seizures. We theorize that a gradual discontinuation of AED treatment allows for a reset of excitatory synaptic strength, preventing seizure onset. In turn, alteration in homeostatic synaptic plasticity may underlie pharmacologically-induced neuropsychiatric effects and recurrent seizures upon discontinuation of AED medication.

## ACKNOWLEDGEMENTS AND CONFLICTS OF INTEREST

This work was supported by the EQUIP Medical Scientist Program, Faculty of Medicine, University of Freiburg (to M.L.), the Berta-Ottenstein Program, Faculty of Medicine, University of Freiburg (to J.S.) and Deutsche Forschungsgemeinschaft (DFG, Project No 520268027 to A.V.). The Galaxy server that was used for some calculations in this study is partially funded by Collaborative Research Centre 992 Medical Epigenetics (DFG grant SFB 992/1 2012) and German Federal Ministry of Education and Research [BMBF grants 031 A538A/A538C RBC, 031L0101B/031L0101C de.NBI-epi, and 031L0106 de.STAIR (de.NBI)].

The authors declare no conflicts of interest.

---

**Supplementary Table 1:**
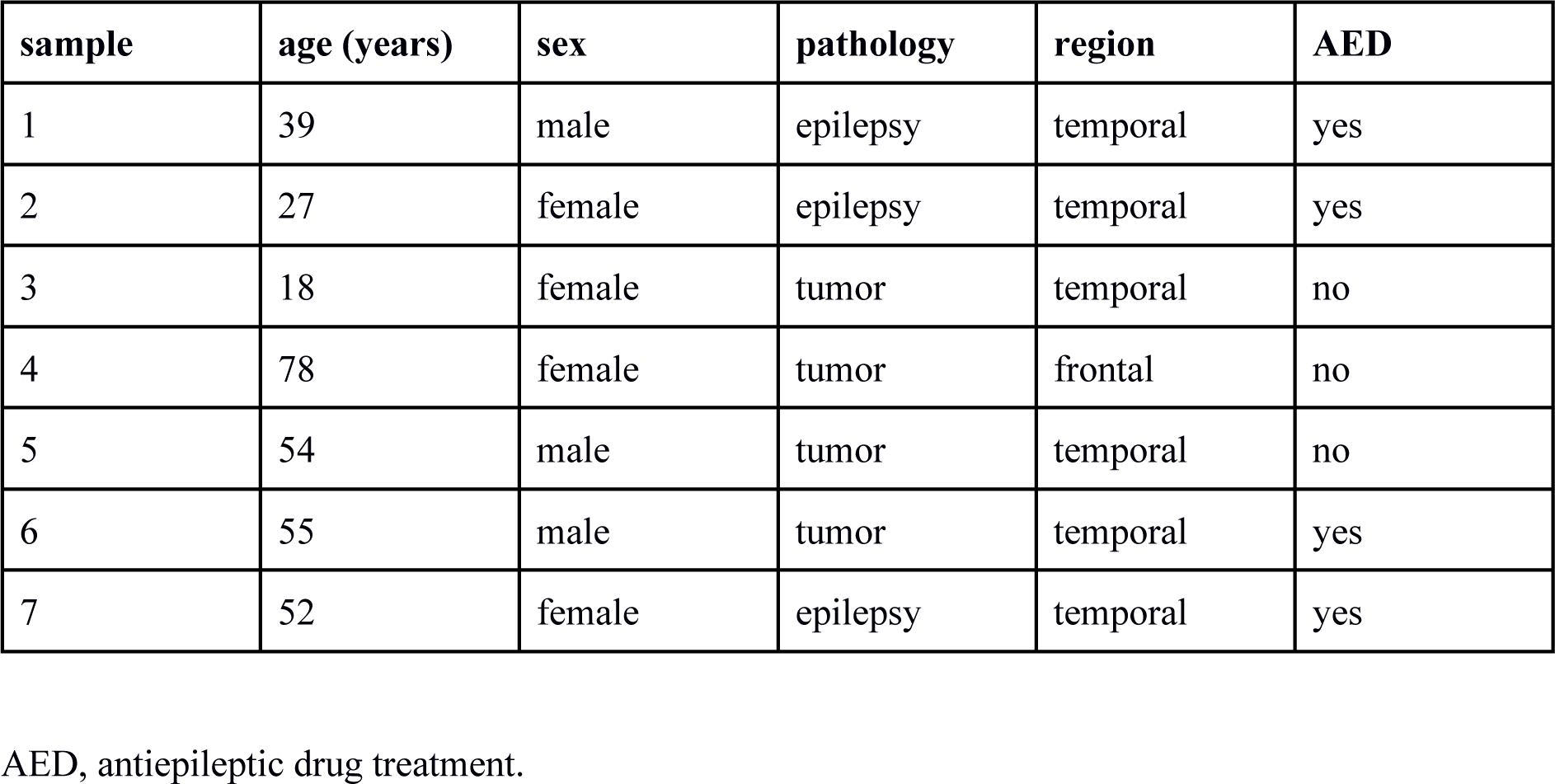
Cortical resection samples.

| sample | age (years) | sex | pathology | region | AED |
| --- | --- | --- | --- | --- | --- |
| 1 | 39 | male | epilepsy | temporal | yes |
| 2 | 27 | female | epilepsy | temporal | yes |
| 3 | 18 | female | tumor | temporal | no |
| 4 | 78 | female | tumor | frontal | no |
| 5 | 54 | male | tumor | temporal | no |
| 6 | 55 | male | tumor | temporal | yes |
| 7 | 52 | female | epilepsy | temporal | yes |

AED, antiepileptic drug treatment.

## Notes

### Competing Interest Statement

The authors have declared no competing interest.

## REFERENCES

Althaus JC, Sutton MA (2021) From mice to men. Elife 10.

Arendt KL, Zhang Y, Jurado S, Malenka RC, Sudhof TC, Chen L (2015) Retinoic Acid and LTP Recruit Postsynaptic AMPA Receptors Using Distinct SNARE-Dependent Mechanisms. Neuron 86:442–456.

Barnes SJ, Sammons RP, Jacobsen RI, Mackie J, Keller GB, Keck T (2015) Subnetwork-Specific Homeostatic Plasticity in Mouse Visual Cortex In Vivo. Neuron 86:1290–1303.

Batut B, van den Beek M, Doyle MA, Soranzo N (2021) RNA-Seq Data Analysis in Galaxy. Methods Mol Biol 2284:367–392.

Chen L, Lau AG, Sarti F (2014) Synaptic retinoic acid signaling and homeostatic synaptic plasticity. Neuropharmacology 78:3–12.

Chen Z, Lusicic A, O’Brien TJ, Velakoulis D, Adams SJ, Kwan P (2016) Psychotic disorders induced by antiepileptic drugs in people with epilepsy. Brain 139:2668–2678.

Davis GW, Bezprozvanny I (2001) Maintaining the stability of neural function: a homeostatic hypothesis. Annu Rev Physiol 63:847–869.

Del Turco D, Deller T (2007) Organotypic entorhino-hippocampal slice cultures--a tool to study the molecular and cellular regulation of axonal regeneration and collateral sprouting in vitro. Methods Mol Biol 399:55–66.

Dubes S, Soula A, Benquet S, Tessier B, Poujol C, Favereaux A, Thoumine O, Letellier M (2022) miR-124-dependent tagging of synapses by synaptopodin enables input-specific homeostatic plasticity. EMBO J 41:e109012.

Eyal G, Verhoog MB, Testa-Silva G, Deitcher Y, Lodder JC, Benavides-Piccione R, Morales J, DeFelipe J, de Kock CP, Mansvelder HD, Segev I (2016) Unique membrane properties and enhanced signal processing in human neocortical neurons. Elife 5.

Gainey MA, Hurvitz-Wolff JR, Lambo ME, Turrigiano GG (2009) Synaptic scaling requires the GluR2 subunit of the AMPA receptor. J Neurosci 29:6479–6489.

Galaxy C (2022) The Galaxy platform for accessible, reproducible and collaborative biomedical analyses: 2022 update. Nucleic Acids Res 50:W345–W351.

Gallinaro JV, Gasparovic N, Rotter S (2022) Homeostatic control of synaptic rewiring in recurrent networks induces the formation of stable memory engrams. PLoS Comput Biol 18:e1009836.

Gidon A, Zolnik TA, Fidzinski P, Bolduan F, Papoutsi A, Poirazi P, Holtkamp M, Vida I, Larkum ME (2020) Dendritic action potentials and computation in human layer 2/3 cortical neurons. Science 367:83–87.

Hunt S et al. (2023) Strong and reliable synaptic communication between pyramidal neurons in adult human cerebral cortex. Cereb Cortex 33:2857–2878.

Kavalali ET, Monteggia LM (2023) Rapid homeostatic plasticity and neuropsychiatric therapeutics. Neuropsychopharmacology 48:54–60.

Keck T, Keller GB, Jacobsen RI, Eysel UT, Bonhoeffer T, Hubener M (2013) Synaptic scaling and homeostatic plasticity in the mouse visual cortex in vivo. Neuron 80:327–334.

Kim J, Tsien RW (2008) Synapse-specific adaptations to inactivity in hippocampal circuits achieve homeostatic gain control while dampening network reverberation. Neuron 58:925–937.

Larkum ME, Wu J, Duverdin SA, Gidon A (2022) The Guide to Dendritic Spikes of the Mammalian Cortex In Vitro and In Vivo. Neuroscience 489:15–33.

Lattanzi S, Trinka E, Del Giovane C, Nardone R, Silvestrini M, Brigo F (2019) Antiepileptic drug monotherapy for epilepsy in the elderly: A systematic review and network meta-analysis. Epilepsia 60:2245–2254.

Lenz M, Galanis C, Kleidonas D, Fellenz M, Deller T, Vlachos A (2019) Denervated mouse dentate granule cells adjust their excitatory but not inhibitory synapses following in vitro entorhinal cortex lesion. Exp Neurol 312:1–9.

Lenz M, Kruse P, Eichler A, Straehle J, Beck J, Deller T, Vlachos A (2021a) All-trans retinoic acid induces synaptic plasticity in human cortical neurons. Elife 10.

Lenz M, Eichler A, Kruse P, Muellerleile J, Deller T, Jedlicka P, Vlachos A (2021b) All-trans retinoic acid induces synaptopodin-dependent metaplasticity in mouse dentate granule cells. Elife 10.

Lenz M, Eichler A, Kruse P, Stohr P, Kleidonas D, Galanis C, Lu H, Vlachos A (2023a) Denervated mouse CA1 pyramidal neurons express homeostatic synaptic plasticity following entorhinal cortex lesion. Front Mol Neurosci 16:1148219.

Lenz M, Eichler A, Kruse P, Galanis C, Kleidonas D, Andrieux G, Boerries M, Jedlicka P, Muller U, Deller T, Vlachos A (2023b) The Amyloid Precursor Protein Regulates Synaptic Transmission at Medial Perforant Path Synapses. J Neurosci 43:5290–5304.

Li J, Park E, Zhong LR, Chen L (2019) Homeostatic synaptic plasticity as a metaplasticity mechanism - a molecular and cellular perspective. Curr Opin Neurobiol 54:44–53.

Lu H, Gallinaro JV, Normann C, Rotter S, Yalcin I (2022) Time Course of Homeostatic Structural Plasticity in Response to Optogenetic Stimulation in Mouse Anterior Cingulate Cortex. Cereb Cortex 32:1574–1592.

Maghsoodi B, Poon MM, Nam CI, Aoto J, Ting P, Chen L (2008) Retinoic acid regulates RARalpha-mediated control of translation in dendritic RNA granules during homeostatic synaptic plasticity. Proc Natl Acad Sci U S A 105:16015–16020.

Maus L, Lee C, Altas B, Sertel SM, Weyand K, Rizzoli SO, Rhee J, Brose N, Imig C, Cooper BH (2020) Ultrastructural Correlates of Presynaptic Functional Heterogeneity in Hippocampal Synapses. Cell Rep 30:3632–3643 e3638.

Moosa ANV (2019) Antiepileptic Drug Treatment of Epilepsy in Children. Continuum (Minneap Minn) 25:381–407.

Park KM, Kim SE, Lee BI (2019) Antiepileptic Drug Therapy in Patients with Drug-Resistant Epilepsy. J Epilepsy Res 9:14–26.

Ratkai A, Tarnok K, Aouad HE, Micska B, Schlett K, Szucs A (2021) Homeostatic plasticity and burst activity are mediated by hyperpolarization-activated cation currents and T-type calcium channels in neuronal cultures. Sci Rep 11:3236.

Raudvere U, Kolberg L, Kuzmin I, Arak T, Adler P, Peterson H, Vilo J (2019) g:Profiler: a web server for functional enrichment analysis and conversions of gene lists (2019 update). Nucleic Acids Res 47:W191–W198.

Reimand J, Isserlin R, Voisin V, Kucera M, Tannus-Lopes C, Rostamianfar A, Wadi L, Meyer M, Wong J, Xu C, Merico D, Bader GD (2019) Pathway enrichment analysis and visualization of omics data using g:Profiler, GSEA, Cytoscape and EnrichmentMap. Nat Protoc 14:482–517.

Rosado J, Bui VD, Haas CA, Beck J, Queisser G, Vlachos A (2022) Calcium modeling of spine apparatus-containing human dendritic spines demonstrates an “all-or-nothing” communication switch between the spine head and dendrite. PLoS Comput Biol 18:e1010069.

Schanzenbacher CT, Sambandan S, Langer JD, Schuman EM (2016) Nascent Proteome Remodeling following Homeostatic Scaling at Hippocampal Synapses. Neuron 92:358–371.

Straehle J, Ravi VM, Heiland DH, Galanis C, Lenz M, Zhang J, Neidert NN, El Rahal A, Vasilikos I, Kellmeyer P, Scheiwe C, Klingler JH, Fung C, Vlachos A, Beck J, Schnell O (2023) Technical report: surgical preparation of human brain tissue for clinical and basic research. Acta Neurochir (Wien) 165:1461–1471.

Sutton MA, Ito HT, Cressy P, Kempf C, Woo JC, Schuman EM (2006) Miniature neurotransmission stabilizes synaptic function via tonic suppression of local dendritic protein synthesis. Cell 125:785–799.

Thapliyal S, Arendt KL, Lau AG, Chen L (2022) Retinoic acid-gated BDNF synthesis in neuronal dendrites drives presynaptic homeostatic plasticity. Elife 11.

Ting JT, Lee BR, Chong P, Soler-Llavina G, Cobbs C, Koch C, Zeng H, Lein E (2018) Preparation of Acute Brain Slices Using an Optimized N-Methyl-D-glucamine Protective Recovery Method. J Vis Exp.

Turrigiano G (2012) Homeostatic synaptic plasticity: local and global mechanisms for stabilizing neuronal function. Cold Spring Harb Perspect Biol 4:a005736.

Turrigiano GG (2008) The self-tuning neuron: synaptic scaling of excitatory synapses. Cell 135:422–435.

Turrigiano GG, Leslie KR, Desai NS, Rutherford LC, Nelson SB (1998) Activity-dependent scaling of quantal amplitude in neocortical neurons. Nature 391:892–896.

van der Meer PB, Dirven L, Fiocco M, Vos MJ, Kouwenhoven MCM, van den Bent MJ, Taphoorn MJB, Koekkoek JAF (2021) First-line antiepileptic drug treatment in glioma patients with epilepsy: Levetiracetam vs valproic acid. Epilepsia 62:1119–1129.

Vercruysse F, Naud R, Sprekeler H (2021) Self-organization of a doubly asynchronous irregular network state for spikes and bursts. PLoS Comput Biol 17:e1009478.

Verhoog MB, Obermayer J, Kortleven CA, Wilbers R, Wester J, Baayen JC, De Kock CPJ, Meredith RM, Mansvelder HD (2016) Layer-specific cholinergic control of human and mouse cortical synaptic plasticity. Nat Commun 7:12826.

Verrotti A, Tambucci R, Di Francesco L, Pavone P, Iapadre G, Altobelli E, Matricardi S, Farello G, Belcastro V (2020) The role of polytherapy in the management of epilepsy: suggestions for rational antiepileptic drug selection. Expert Rev Neurother 20:167–173.

Vlachos A, Ikenberg B, Lenz M, Becker D, Reifenberg K, Bas-Orth C, Deller T (2013) Synaptopodin regulates denervation-induced homeostatic synaptic plasticity. Proc Natl Acad Sci U S A 110:8242–8247.

Wang J, Huang P, Song Z (2019a) Comparison of the relapse rates in seizure-free patients in whom antiepileptic therapy was discontinued and those in whom the therapy was continued: A meta-analysis. Epilepsy Behav 101:106577.

Wang X, He R, Zheng R, Ding S, Wang Y, Li X, Hua Y, Zeng Q, Xia N, Zhu Z, Kwan P, Xu H (2019b) Relative Seizure Relapse Risks Associated with Antiepileptic Drug Withdrawal After Different Seizure-Free Periods in Adults with Focal Epilepsy: A Prospective, Controlled Follow-Up Study. CNS Drugs 33:1121–1132.

Wen W, Turrigiano GG (2021) Developmental Regulation of Homeostatic Plasticity in Mouse Primary Visual Cortex. J Neurosci 41:9891–9905.

Wu CH, Ramos R, Katz DB, Turrigiano GG (2021) Homeostatic synaptic scaling establishes the specificity of an associative memory. Curr Biol 31:2274–2285 e2275.

Wu CH, Tatavarty V, Jean Beltran PM, Guerrero AA, Keshishian H, Krug K, MacMullan MA, Li L, Carr SA, Cottrell JR, Turrigiano GG (2022) A bidirectional switch in the Shank3 phosphorylation state biases synapses toward up- or downscaling. Elife 11.

Yee AX, Hsu YT, Chen L (2017) A metaplasticity view of the interaction between homeostatic and Hebbian plasticity. Philos Trans R Soc Lond B Biol Sci 372.

