## Supplementary figures and images for "Antiepileptic medication strengthens excitatory neurotransmission in pyramidal neurons of the adult human neocortex"

### Figure S1

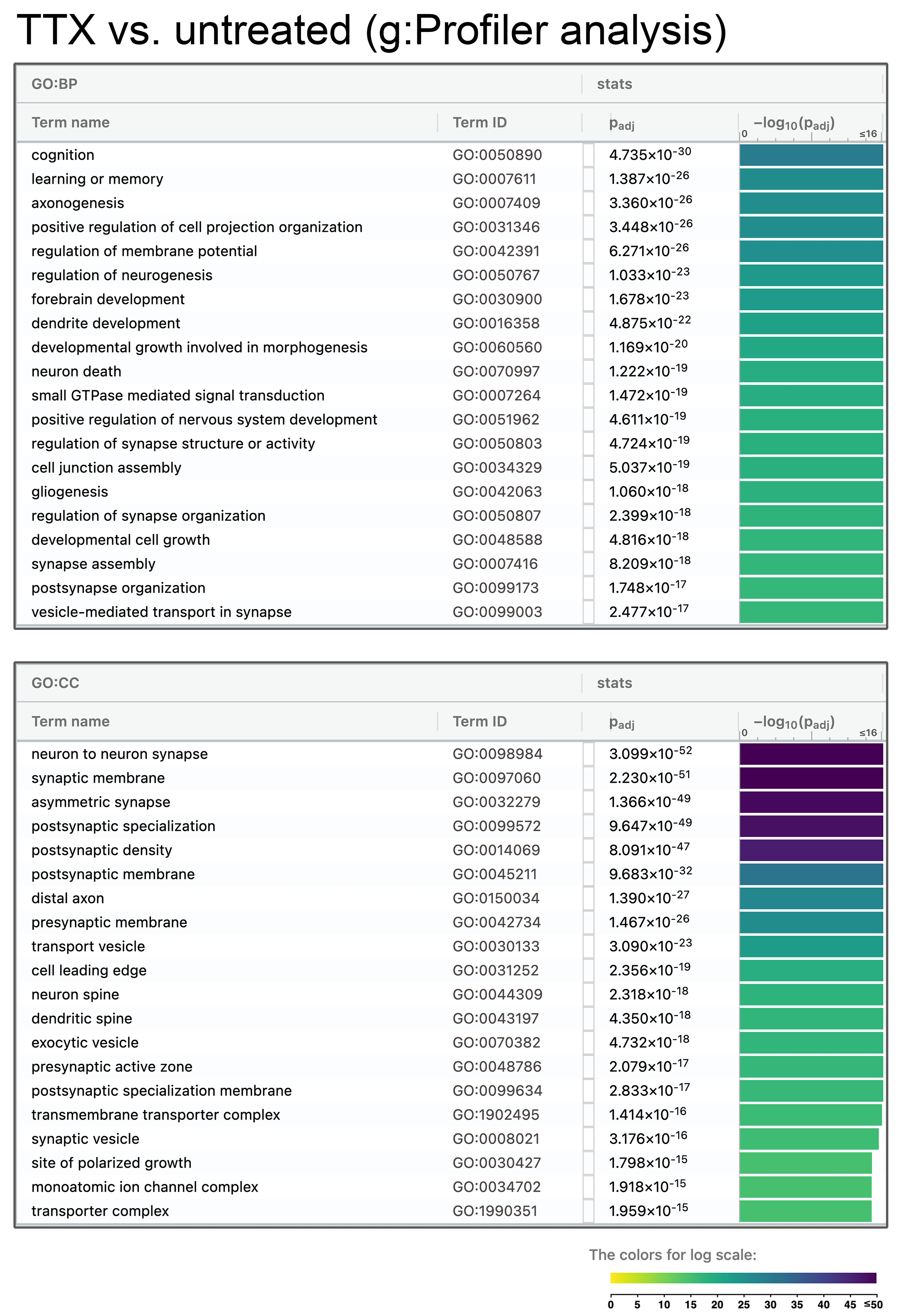

### Figure S2

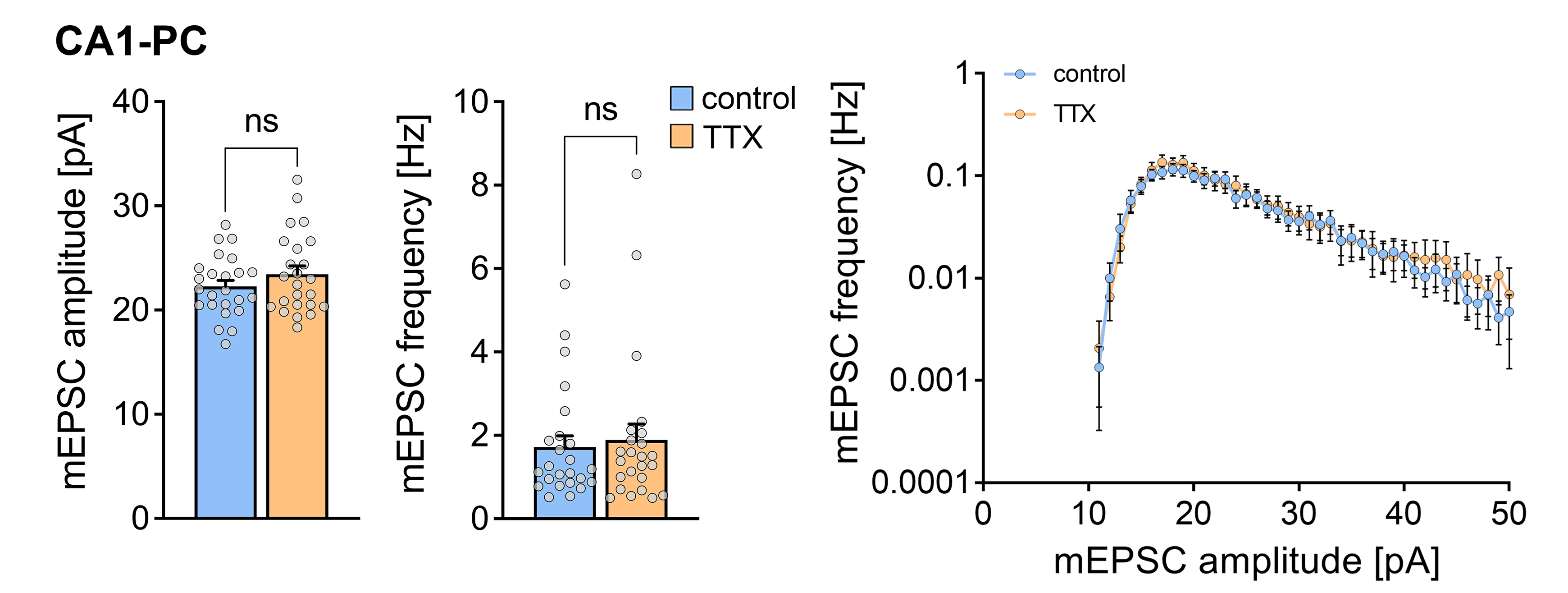

### Figure S3

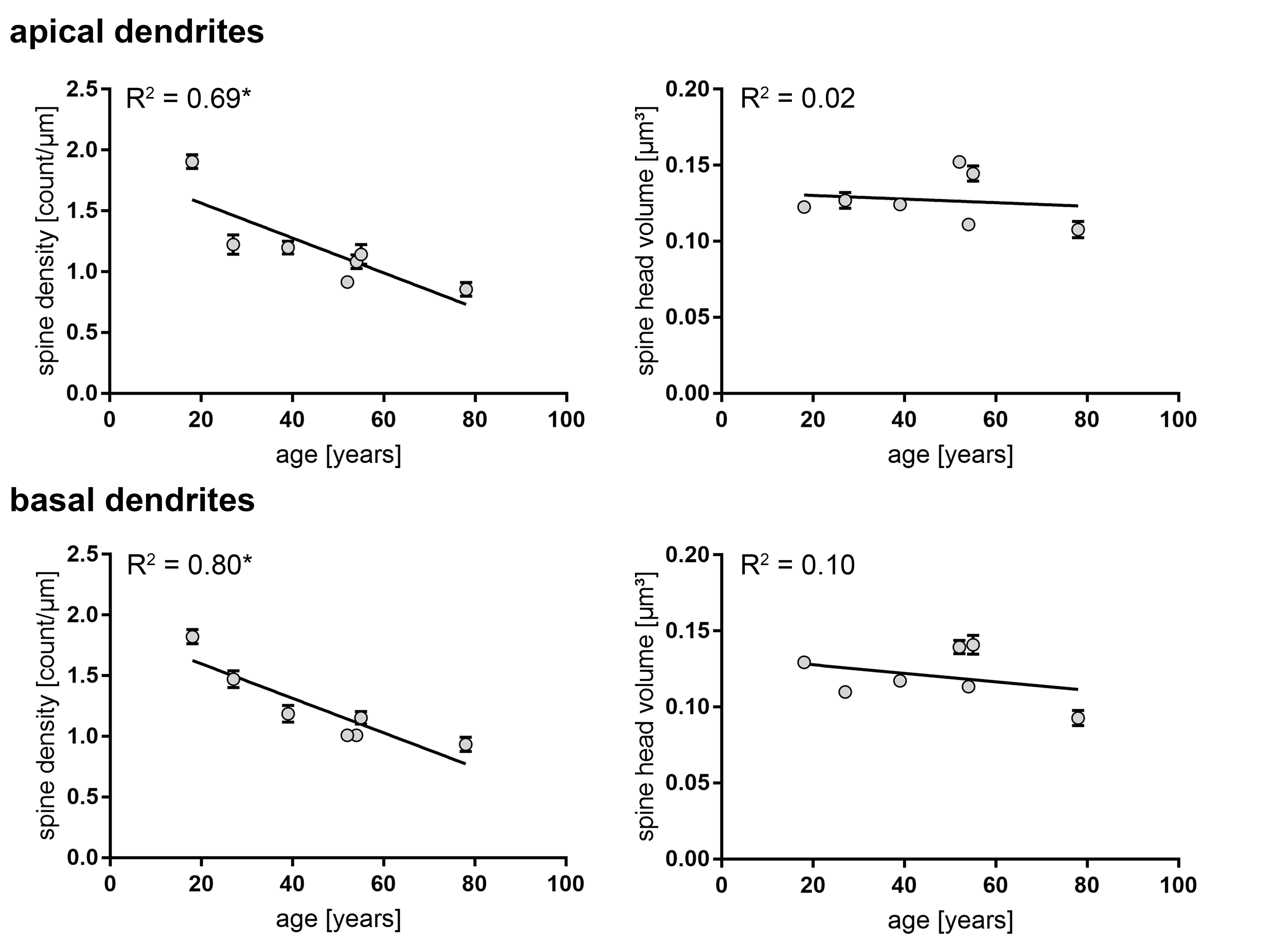
